# Multimodal Structural and Regulatory Architecture of Coding and Non-Coding Variants Across the Orofacial Clefting *TP63–IRF6–GRHL3* Network

**DOI:** 10.64898/2026.11.17.767356

**Authors:** Filipe Martins Vono, Valeria Gonzaga Santos, Andre Naufel de Toledo

**Affiliations:** Centro Universitário de Belo Horizonte (UniBH), Belo Horizonte, Minas Gerais, Brazil; American Global Tech University, Orlando, Florida, USA

## Abstract

Pathogenic variation across the *TP63*–*IRF6*–*GRHL3* transcriptional network causes syndromic and nonsyndromic orofacial clefts (OFCs), yet scalar sequence predictors collapse distinct failure modes and leave isoform-specific coding variants, splice-altering indels, and distal enhancer alleles unresolved. Here, we integrate 29 AlphaFold 3 (AF3) assemblies (10 wild-type, 19 mutant), thermodynamic stability (ΔΔG) calculations, and AlphaGenome regulatory modeling across clinical, saturation-mutagenesis, and reporter benchmarks. In strict leave-one-gene-out (LOGO) coding benchmarks, replacing paralog crystal templates with full-length human AF3 structures improves *IRF6* classification (biophysical AUROC rising from 0.7844 to 0.8967; ΔΔG from 0.6461 to 0.8516) and resolves canonical isoform gaps in the 602-residue MANE Select GRHL3 homodimer. Projecting *IRF6* missense variants onto a two-dimensional stability-versus-interface phase plane separates dominant-negative popliteal pterygium syndrome (PPS) poisons from haploinsufficient Van der Woude syndrome (VWS1) alleles (VWS1-versus-PPS Quadrant I odds ratio [OR] = 0.0263 [reciprocal OR = 38.0], *p* = 4.652 × 10^−3^) and distinguishes DNA-contact poisons from dimer-unfolding alleles across 12 zebrafish *in vivo* assays. Atomistic base-readout contacts in the AF3 TP63–*MCS9.7* tetramer mirror loss-of-function profiles in MaveDB (ρ = 0.6048, *p* = 6.52 × 10^−4^) and AlphaGenome keratinocyte DNase predictions (ρ = 0.6040, *p* = 6.65 × 10^−4^). Across non-coding benchmarks, AlphaGenome splicing tracks reach an AUROC of 0.9876 while prioritizing cryptic splice variants of uncertain significance, and epithelial chromatin tracks capture enhancer grammar, plasmid-cloning artifacts, and context-dependent *ARHGAP29, FOXE1*, and local *8q24* chromatin effects within AlphaGenome’s 1.049-Mb horizon.

## Introduction

Craniofacial morphogenesis and palatal fusion require spatiotemporal coordination between basal ectodermal progenitors and the superficial oral periderm, governed by a hierarchical transcriptional network linking *TP63, IRF6*, and *GRHL3* (1–4). Heterozygous germline variants across these loci cause an overlapping yet mechanistically diverse spectrum of peridermopathies and orofacial clefts (OFCs). Coding mutations in *TP63* cause ectrodactyly, ectodermal dysplasia, and cleft lip/palate (EEC), ankyloblepharon-ectodermal defects-cleft lip/palate (AEC), and Rapp-Hodgkin (RHS) syndromes through domain-specific perturbations of DNA binding, tetramerization, or sterile alpha motif (SAM) folding (1, 5–7). Mutations in *IRF6* account for ∼70% of Van der Woude syndrome type 1 (VWS1) cases and cause the more severe popliteal pterygium syndrome (PPS) (2, 8, 9), while downstream variants in *GRHL3* cause Van der Woude syndrome type 2 (VWS2) and confer isolated cleft-palate risk across diverse populations (4, 10–14).

Susceptibility across this network extends seamlessly into *cis*-regulatory architecture. TP63 activates *IRF6* in oral epithelium via *MCS9*.*7*, a conserved upstream enhancer where common risk alleles (rs642961) and rare mutations disrupt craniofacial activity (3, 15, 16). Genome-wide association studies (GWASs), targeted sequencing, and massively parallel reporter assays (MPRAs) further show that non-coding variants near *IRF6, TP63, 8q24*/*MYC, FOXE1, MAFB*, and *1p22* (*ABCA4*/*ARHGAP29*) drive a major share of nonsyndromic OFC heritability (17–22).

Despite progress in deep learning for human genetics, two interpretive barriers persist. First, evolutionary missense predictors such as AlphaMissense achieve strong binary sorting (23) yet collapse distinct biophysical mechanisms into a scalar score. Although most pathogenic missense alleles act via thermodynamic destabilization (24), clinical severity in oligomeric transcription factors hinges on whether a variant triggers haploinsufficient degradation or preserves a folded subunit that poisons DNA binding or oligomeric switches—a distinction scalar scores cannot resolve across *IRF6* and *TP63* (5, 8, 9, 25). Moreover, experimental crystal templates remain fragmented across isolated domains or paralogs, while canonical transcript choices omit segments such as the C-terminal dimerization tail of 602-residue MANE Select *GRHL3*. Second, short-window non-coding motif models miss insertion-deletion (indel) splice junctions, conflate episomal plasmid-cloning artifacts with enhancer loss of function, and lack the sub-megabase context needed to link distal OFC risk alleles to effector genes.

Here, we integrate 29 AlphaFold 3 (AF3) protein–DNA, Zn^2+^-coordinated, and phosphoswitch assemblies (26) with ThermoMPNN ΔΔG calculations (27) and AlphaGenome regulatory modeling (28) to evaluate zero-shot biophysical generalization, resolve isoform gaps, separate dominant-negative PPS from haploinsufficient VWS1 alleles, predict zebrafish periderm rupture (8, 25), and bridge atomic TP63–*MCS9*.*7* readout to clinical splicing, enhancer saturation mutagenesis (29), and GWAS effector regulation (22).

## Material and Methods

### Coding Variant Cohorts and Structural Templates

Pathogenic/likely pathogenic missense variants in *TP63, IRF6*, and *GRHL3* from ClinVar and syndromic OFC cohorts were paired against gnomAD (GRCh38) missense controls at AC ≥ 2, with AC ≥ 5 and AC ≥ 10 sweeps for incomplete-penetrance *GRHL3* alleles (10, 11). Crystal benchmarks used human TP63 DBD (PDB: 3QYN) (30, 31), OD (4A9Z) (32), and SAM (2Y9U) (6); paralog *IRF6* templates from murine Irf1 DBD (1IF1) (33) and dimeric human IRF5 IAD (3DSH) (34); and human GRHL3 DBD (5MPF) (35). Domain benchmarks comprised Domainome 1.0 abundance DMS on TP63 SAM 2Y9U (*n* = 1,244: 1,198 DMS, 27 gnomAD AC ≥ 2, 19 ClinVar AEC/RHS pathogenic/likely pathogenic variants) (24), 37 stratified TP63 DBD mutations on 3QYN (5), and 893 saturation variants on TP63 OD homotetramer 4A9Z alongside ClinVar VUS and benign/likely benign alleles (32). Full-length human AF3 assemblies expanded crystal-mapped clinical cohorts across linkers, human IAD loops, and 602-residue MANE Select *GRHL3* (NM_198173.3) (**Table 1**). Syndrome-resolved *IRF6* alleles (6 PPS, 20 VWS1) and 12 zebrafish-assayed *IRF6* variants were compiled from UniProt and *in vivo* studies (8, 25). Precomputed AlphaMissense scores (23) restrict *GRHL3* to residues 1–565.

**Table 1.**
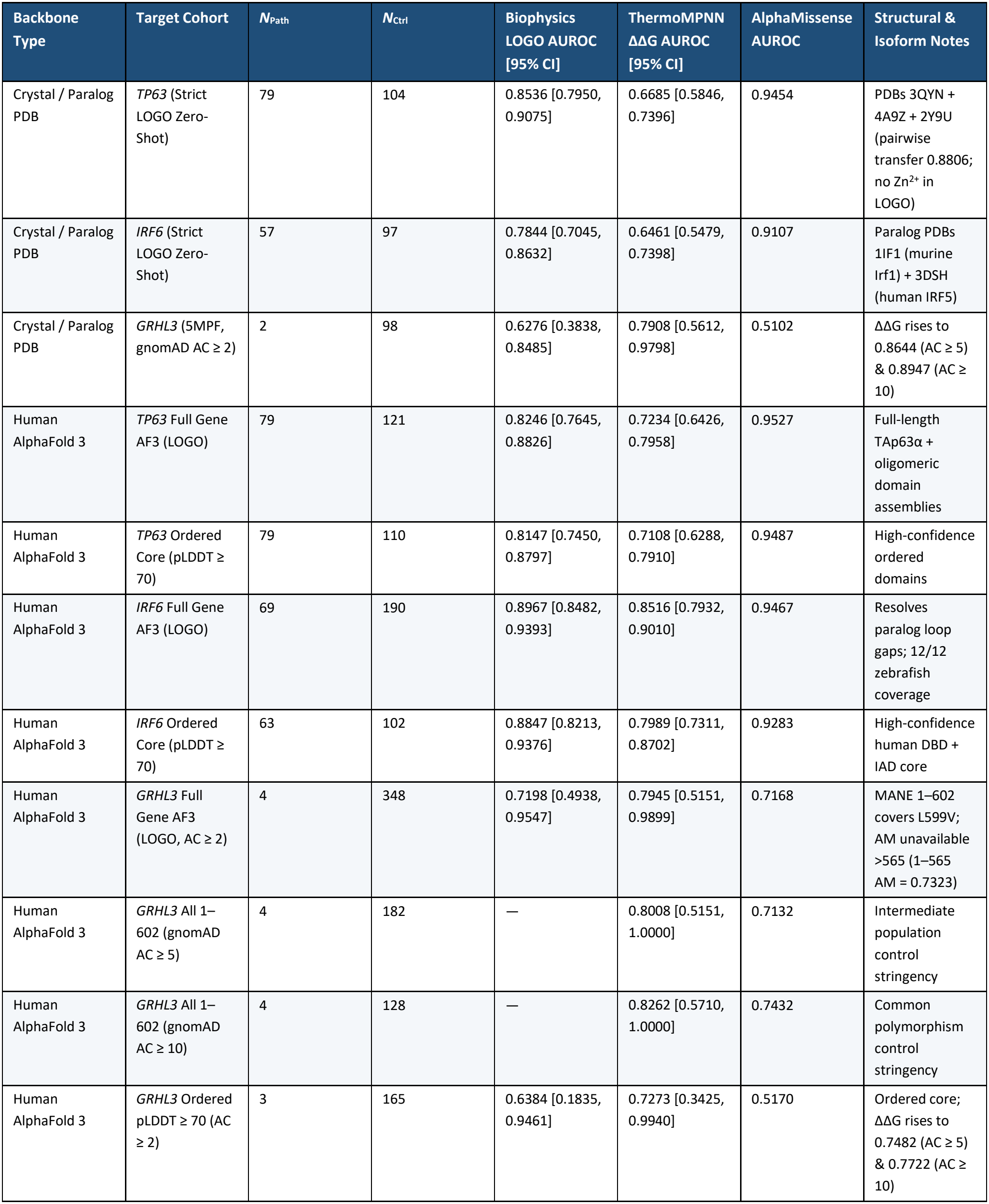
Coding Missense Pathogenicity Benchmarks Across Experimental Crystal/Paralog Templates and Full-Length Human AlphaFold 3 Structures.

### AlphaFold 3 Modeling and Biophysical Feature Extraction

Twenty-nine assemblies (10 wild-type [WT], 19 mutant; **Table 2**; **Table S6**) were predicted with AlphaFold 3 via AlphaFold Server under non-commercial Output Terms of Use (26). WT runs comprised Zn^2+^-bound full-length TAp63α monomer, TP63 OD tetramer, TP63 SAM monomer, and Zn^2+^-coordinated TP63 DBD tetramers on palindromic DNA or native 28-bp *MCS9*.*7* (fold_wt_tp63_dbd_tet_mcs97_zn); apo full-length IRF6 (fold_wt_irf6_fl_mon), apo IRF6 IAD monomer, ISRE-bound IRF6 DBD dimer, and C-terminally phosphorylated IRF6 IAD dimer (fold_wt_irf6_iad_phospho_dim); and DNA-bound 602-residue MANE Select GRHL3 homodimer (fold_wt_grhl3_mane602_dim_dna). No PRKCI kinase–IRF6 complex was modeled. Extracted metrics included Chain A mean pLDDT, pTM, ipTM, sub-block prot_prot_iptm_max and prot_dna_iptm_max, minimum inter-chain PAE (prot_prot_pae_min, prot_dna_pae_min), DNA-centered Cα RMSD versus WT, and C3H1 Zn^2+^ bond/angle geometry. ThermoMPNN (27) ΔΔG (kcal/mol) was computed across domain-matched (ΔΔG_pred_), full-length apo monomer (ΔΔG_AF3,FL_), and phosphorylated dimer (ΔΔG_AF3,pdim_) backbones alongside heavy-atom distances to DNA bases (d_DNA,base_), phosphates (d_DNA,PO4_), overall DNA (d_DNA_), oligomeric partners (d_olig_), and Zn^2+^, buried interface area (ΔSASA_int_), relative solvent accessibility (RSA), and per-base-pair *MCS9*.*7* hydrogen bonds (≤ 3.5 Å).

**Table 2.** Representative AlphaFold 3 Biomolecular Assemblies, Protein–DNA/Subunit Interface Metrics, and Zn^2+^ Coordination Perturbations (Subset of 29 Completed Runs)

| AF3 Job Name | Chains (Prot / DNA) | Chain A Mean pLDDT | pTM | ipTM | Max Prot–Prot ipTM | Max Prot–DNA ipTM | Min Prot–Prot PAE (Å) | Min Prot–DNA PAE (Å) | Zn<sup>2+</sup> Max Bond (Å) / Min Angle (°) | Primary Mechanistic Readout |
|---|---|---|---|---|---|---|---|---|---|---|
| fold_wt_tp63_dbd_tet_mcs97_zn | 4 / 2 | 93.10 | 0.88 | 0.85 | 0.85 | 0.63 | 1.34 | 4.07 | 2.57 / 99.32 | Native *MCS9.7* DNA tetramer; intact tetrahedral Zn<sup>2+</sup> |
| fold_wt_tp63_dbd_tet_paldna_zn | 4 / 2 | 92.11 | 0.78 | 0.74 | 0.81 | 0.54 | 1.50 | 5.92 | 2.56 / 102.55 | Lower DNA interface confidence than native *MCS9.7* |
| fold_mut_tp63_dbd_c312y_mcs97 | 4 / 2 | 93.25 | 0.87 | 0.85 | 0.85 | 0.62 | 1.47 | 4.26 | **3.97 / 15.53** | Catastrophic Zn<sup>2+</sup> coordination sphere collapse |
| fold_mut_tp63_dbd_r243w_mcs97 | 4 / 2 | 91.13 | 0.87 | 0.84 | 0.84 | **0.58** | 1.47 | **4.93** | 2.55 / 92.27 | Reduced DNA ipTM, widened DNA PAE, Zn<sup>2+</sup>-loop strain |
| fold_mut_tp63_dbd_r318h_mcs97 | 4 / 2 | 93.35 | 0.88 | 0.85 | 0.85 | 0.61 | 1.32 | 4.60 | 2.56 / 99.76 | Mild major-groove PAE expansion |
| fold_wt_tp63_sam_mon | 1 / 0 | 86.66 | 0.72 | — | — | — | — | — | — | Ordered SAM domain (L562P drops pLDDT to 81.50, pTM 0.67) |
| fold_wt_irf6_iad_phospho_dim | 2 / 0 | 82.97 | 0.52 | 0.60 | 0.60 | — | 2.77 | — | — | Active phosphorylated IRF6 IAD homodimer |
| fold_mut_irf6_iad_s424l_pdim | 2 / 0 | 80.83 | 0.52 | **0.58** | **0.58** | — | **3.21** | — | — | PPS phosphoswitch mutation weakens IAD dimer docking |
| fold_mut_irf6_iad_v274i_pdim | 2 / 0 | 82.52 | 0.54 | 0.61 | 0.61 | — | 2.64 | — | — | Rescued benign control preserves tight IAD dimerization |
| fold_wt_irf6_dbd_dim_isre_dna | 2 / 2 | 82.80 | 0.40 | 0.23 | 0.05 | 0.19 | 25.72 | 15.07 | — | Isolated DBD lacks tight clamping (R84C DNA ipTM = 0.15) |
| fold_wt_grhl3_mane602_dim_dna | 2 / 2 | 62.29 | 0.46 | 0.45 | 0.43 | 0.75 | 2.13 | 3.88 | — | Confident DNA-bound GRHL3 MANE 602-aa homodimer |
| fold_mut_grhl3_l599v_dim_dna | 2 / 2 | 63.87 | 0.47 | 0.46 | 0.44 | 0.73 | 2.08 | 3.82 | — | C-terminal MANE dimer mutant (ΔΔG = 1.11 kcal/mol) |

*Note*: Complete metrics for all 29 completed AlphaFold 3 runs (10 WT + 19 Mutants) are provided in **Table S6**. All structures predicted with AlphaFold Server; use is subject to the AlphaFold Server Output Terms of Use.

### Zero-Shot Classification and Statistical Analysis

Regularized logistic classifiers combining ThermoMPNN ΔΔG, RSA, and interface distances were evaluated via strict three-gene LOGO and pairwise cross-gene transfer (excluding Zn^2+^ across genes), 5-fold codon-grouped cross-validation within *TP63* and *IRF6*, and ordered-core (pLDDT ≥ 70) filtering, using 1,000-iteration stratified bootstrap 95% CIs and two-sided DeLong tests (36). Quadrant I *IRF6* poisons were defined *a priori* as ΔΔG_pred_ < 0.75 kcal/mol with d_DNA_ ≤ 3.5 Å or C-terminal phosphoswitch lid (CTD_Phospho) localization and tested by two-sided Fisher’s exact test.

### Non-Coding Benchmarks Across AlphaGenome Modalities

AlphaGenome GRCh38 predictions (28) (non-commercial terms) were stratified into observed, imputed_supported, and imputed_exploratory (gingival epithelial CAGE includes observed and imputed_exploratory tracks; DNASE and RNA_SEQ are imputed_exploratory) across three tiers (**Table 3**): (1) **Tier NC-1**, testing 23 ClinVar pathogenic/likely pathogenic variants against gnomAD controls across *IRF6, TP63*, and *GRHL3* (*N* = 357 [23 pathogenic, 334 controls] for SPLICE_JUNCTIONS, SPLICE_SITE_USAGE, and SPLICE_SITES, excluding 3 upstream controls outside splice-annotated transcripts; *N* = 360 [23 pathogenic, 337 controls] for RNA_SEQ and MaxEntScan |Δscore| [37]; *N* = 23 [20 pathogenic, 3 controls] for precomputed CADD PHRED SNVs) plus 101 ClinVar VUS; (2) **Tier NC-2**, spanning all *N* = 1,785 SNVs across 601-bp *IRF6 MCS9*.*7* (chr1:209,815,790– 209,816,390; 620 SNVs at *p* < 0.01; 364 LoF; 42 severe LoF |score| ≥ 1.0) versus keratinocyte MaveDB MPRA scores (29), regressing imputed JUND (sk_jund_imp) onto observed skin keratinocyte DNASE; and (3) **Tier NC-3**, evaluating *N* = 65 OFC MPRA SNVs (*q* < 0.05; *N* = 62 for gene-mapped RNA_SEQ), *n* = 13 luciferase SNVs (22), and 16-KB to 1-MB (1,048,576-bp) context scans at *8q24*/*MYC* (rs987525), *1p22*/*ABCA4*–*ARHGAP29* (rs560426), *FOXE1* (rs10984103), and *IRF6* (rs661849).

**Table 3.**
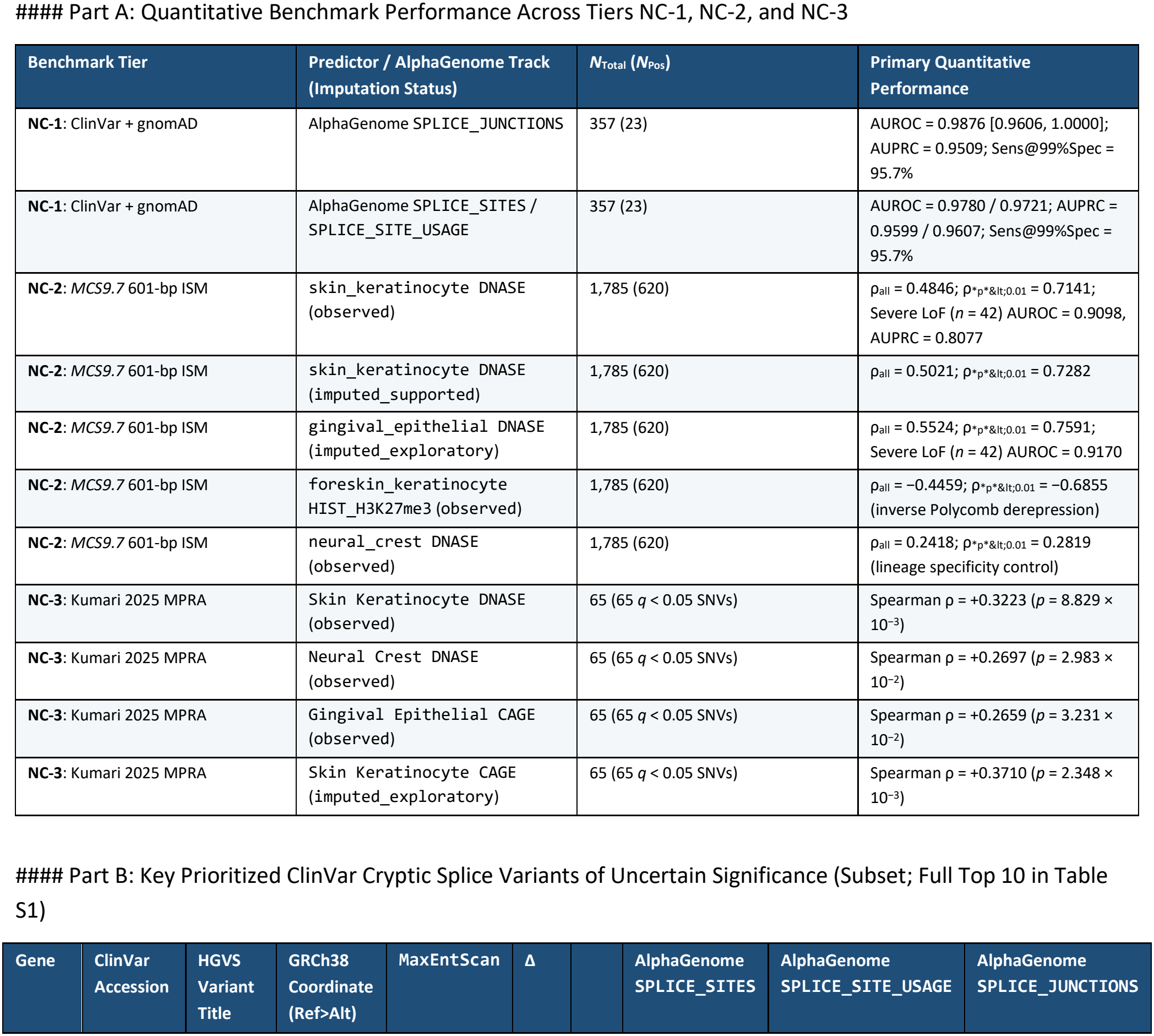
Non-Coding Multimodal Validation Summary (Tiers NC-1, NC-2, NC-3) and Top Prioritized ClinVar Cryptic Splice VUS.

## Results

### Full-Length Human AlphaFold 3 Assemblies Resolve Paralog Template and Isoform Gaps in Zero-Shot Coding Benchmarks

We first evaluated whether explicit three-dimensional biophysical features—thermodynamic destabilization (ΔΔG), solvent burial, and proximity to DNA, oligomeric interfaces, or Zn^2+^—generalize across *TP63, IRF6*, and *GRHL3* without gene-specific supervision (**Figure 1A–D**; **Table 1**; **Table S5**). On crystal templates, strict three-gene LOGO biophysical classification achieved an AUROC of 0.8536 on *TP63* (*P* = 79, *C* = 104)—and 0.8806 under pairwise *IRF6*-to-*TP63* transfer (**Figure 1D**)—outperforming ThermoMPNN ΔΔG alone (0.6685; DeLong *p* = 1.657 × 10^−7^). Domain benchmarks explained this gain: in the globular TP63 SAM domain (2Y9U), ΔΔG tracked Domainome 1.0 cellular abundance across *n* = 1,244 variants (ρ = −0.638, *r* = −0.755; including 19 ClinVar AEC/RHS unfolders such as G569R, F565V, and I549N) (24) (**Figure 1A**), reaching an AUROC of 0.9493 on clinical SAM alleles (0.9240 for 5-fold codon-grouped biophysics). Conversely, across 37 stratified TP63 DBD mutations on 3QYN (5), DNA-contact (R318H, R343W) and Zn^2+^ (C312Y) alleles preserved monomer stability while clustering near DNA/Zn^2+^ (**Figure 1B**); monomer ΔΔG dropped to 0.5691 on clinical DBD alleles (*P* = 56, *C* = 53), whereas 3D interface geometry restored 5-fold codon-grouped AUROC to 0.8814 (0.8918 across full crystal *TP63* with Zn^2+^). Comparing homotetramer versus monomer ΔΔG across 893 saturation variants in the TP63 oligomerization domain (4A9Z) (32) similarly unmasked ClinVar tetramer-interface VUS Y402H (0.38 → 1.75 kcal/mol; ΔΔΔG = +1.37 kcal/mol) and E422K (0.09 → 1.31 kcal/mol; +1.22 kcal/mol) alongside P428L, while keeping benign T434M neutral (**Figure 1C**).

**Figure 1.**
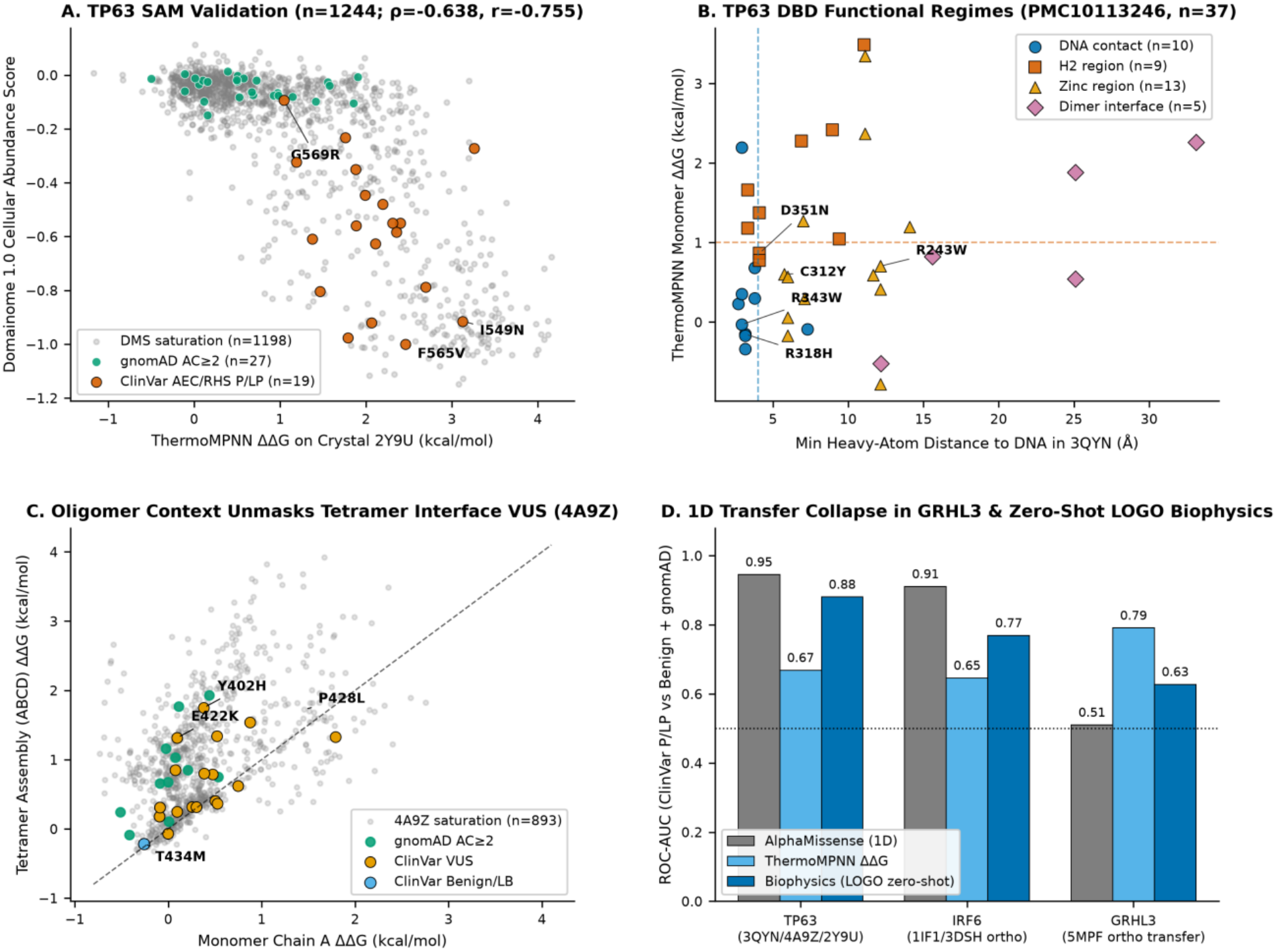
Crystal-structure biophysical validation across TP63 domains, oligomeric VUS unmasking, and cross-gene zero-shot coding benchmarks. (**A**) TP63 SAM domain validation (*n* = 1,244; Spearman ρ = −0.638, Pearson *r* = −0.755) plotting Domainome 1.0 cellular abundance scores against ThermoMPNN ΔΔG (kcal/mol) on crystal structure 2Y9U across DMS saturation variants (*n* = 1,198), gnomAD AC ≥ 2 controls (*n* = 27), and ClinVar AEC/RHS pathogenic/likely pathogenic alleles (*n* = 19, highlighting G569R, F565V, and I549N). (**B**) TP63 DBD functional regimes (*n* = 37) (5) plotting ThermoMPNN monomer ΔΔG against minimum heavy-atom distance to DNA (Å) in crystal structure 3QYN across DNA-contact (*n* = 10; R318H, R343W), helix H2 (*n* = 9; D351N), Zn^2+^-coordinating (*n* = 13; C312Y), and dimer-interface (*n* = 5; R243W) mutations. (**C**) Oligomeric context unmasks tetramer-interface VUS on crystal structure 4A9Z, comparing homotetramer assembly (ABCD) ΔΔG against monomer (Chain A) ΔΔG across 4A9Z saturation variants (*n* = 893), gnomAD AC ≥ 2 controls, ClinVar VUS, and ClinVar benign/likely benign alleles, highlighting unmasked interface VUS Y402H (monomer 0.38 → tetramer 1.75 kcal/mol; ΔΔΔG = +1.37 kcal/mol) and E422K (0.09 → 1.31 kcal/mol; ΔΔΔG = +1.22 kcal/mol) alongside P428L and benign T434M. (**D**) 1D transfer collapse in *GRHL3* and zero-shot biophysical generalization, comparing ROC-AUC for AlphaMissense (1D), ThermoMPNN ΔΔG, and zero-shot Biophysics across *TP63* (3QYN/4A9Z/2Y9U: 0.95 / 0.67 / 0.88 [pairwise transfer; 0.85 three-gene LOGO]), *IRF6* (1IF1/3DSH paralog templates: 0.91 / 0.65 / 0.77 [pairwise transfer; 0.78 three-gene LOGO]), and *GRHL3* (5MPF transfer: 0.51 / 0.79 / 0.63).

On *IRF6* (*P* = 57, *C* = 97), chimeric murine Irf1 (1IF1) and human IRF5 (3DSH) paralog crystals constrained LOGO biophysical AUROC to 0.7844 (pairwise 0.77) and ΔΔG to 0.6461 (**Figure 1D**). Transitioning to full-length human AF3 structures expanded *IRF6* coverage by 68.2% (259 variants: 69 pathogenic, 190 controls) and raised strict LOGO biophysical AUROC to 0.8967 (ordered core 0.8847) and ΔΔG AUROC to 0.8516, approaching supervised AlphaMissense (0.9467) (**Figure 4D**; **Table 1**).

Modeling the DNA-bound 602-residue MANE Select GRHL3 homodimer (fold_wt_grhl3_mane602_dim_dna) resolved two annotation gaps (**Figure 1D**; **Figure 4D**; **Table 1**). First, precomputed AlphaMissense tables terminate at residue 565, omitting pathogenic C-terminal dimerization allele L599V (AF3 ΔΔG_pred_ = 1.1087 kcal/mol, ΔΔG_dim_ = 1.0713 kcal/mol, d_olig_ = 7.87 Å). Second, AlphaMissense scores pathogenic subunit-interface allele R298H as benign (0.1382), whereas the AF3 homodimer buries Arg298 across the inter-monomer cleft (d_olig_ = 2.65 Å, ΔSASA_int_ = 120.35 Å^2^). Although sparse Mendelian positives (*P* = 4 full gene, *P* = 3 ordered core, *P* = 2 on 5MPF) yield wide bootstrap CIs (5MPF DeLong *p* = 0.2943; **Table 1**) that preclude powered superiority claims, point-estimate ThermoMPNN ΔΔG rankings consistently exceeded AlphaMissense across AC ≥ 2, ≥ 5, and ≥ 10 on full 1–602 AF3 (0.7945–0.8262 vs 0.7132–0.7432; unmasked residues 1–565 [*P* = 3, *C* = 348] AlphaMissense = 0.7323), ordered cores, and crystal 5MPF (**Table 1**).

### A Two-Dimensional Biophysical Phase Plane Separates Dominant-Negative PPS from Haploinsufficient VWS1 and Predicts *In Vivo* Embryonic Rupture

Despite strong binary sorting on *IRF6* (AUROC = 0.9467), AlphaMissense saturated near unity across both dominant-negative PPS (median = 0.9989; R84C = 0.9998, S424L = 0.9899) and haploinsufficient VWS1 (median = 0.9897; R84G = 0.9999, T64I = 0.9995) (**Figure 4B**; **Table S5**). Projecting 26 syndrome-resolved *IRF6* missense alleles (6 PPS, 20 VWS1) onto a two-dimensional human AF3 phase plane—pairing domain-matched monomer ΔΔG_pred_ against direct DNA proximity (d_DNA_ ≤ 3.5 Å) or C-terminal phosphoswitch lid (CTD_Phospho) localization— isolated Quadrant I folded poisons (ΔΔG_pred_ < 0.75 kcal/mol): 4/6 PPS alleles (R84C −0.3423, R84L −0.3353, and R84H +0.3089 kcal/mol at d_DNA_ = 2.49 Å; S424L +0.0865 kcal/mol) versus 1/20 VWS1 alleles (N88H, +0.2302 kcal/mol, d_DNA_ = 3.39 Å, d_DNA,base_ = 4.598 Å; Fisher’s exact VWS1-versus-PPS OR = 0.0263 [reciprocal PPS enrichment OR = 38.0], *p* = 4.652 × 10^−3^). At codon Arg84 itself, helix-destabilizing R84G (ΔΔG_pred_ = +1.1351 kcal/mol) shifted out of Quadrant I into haploinsufficient VWS1, whereas incomplete paralog crystals (6 PPS, 8 VWS1) failed to reach significance (OR = 0.07, *p* = 9.091 × 10^−2^).

Across 12 human *IRF6* variants assayed in zebrafish embryonic periderm (8, 25), human AF3 restored 12/12 (100%) coverage versus 7/12 (58.3%) WT-identical paralog crystal sites (**Figure 4B**,**D**; **Table S5**). Evaluated on full-length apo monomer fold_wt_irf6_fl_mon (ΔΔG_AF3,FL_) and phosphorylated IAD dimer fold_wt_irf6_iad_phospho_dim (ΔΔG_AF3,pdim_), all four lethal-rupture alleles bifurcated into hyper-stable DNA poisons (K66T: ΔΔG_AF3,FL_ = −0.7606 kcal/mol, d_DNA_ = 1.86 Å; R84C: −0.5124 kcal/mol, d_DNA,base_ = 2.49 Å) or severe IAD dimer-core unfolders (L294P: +2.5269 / +2.6942 kcal/mol; G325E: +2.3997 / +2.4415 kcal/mol). All eight mRNA-rescued alleles (R45Q, R45W, G70R, V274I, R400Q, R9W, N88H, T100A) exhibited mild-to-moderate ΔΔG_AF3,FL_ (−0.2384 to +1.6021 kcal/mol) without close nucleobase contact (d_DNA,base_ ≥ 4.598 Å), consistent with dosage-surmountable hypomorphs despite near-unity AlphaMissense scores (0.9911–0.9975).

### Atomistic AlphaFold 3 Protein–DNA Readout Bridges *TP63* Coding Mechanisms to *MCS9*.*7* Enhancer Vulnerability

Across all 29 AF3 assemblies (**Figure 4A,C**; **Table 2**; **Table S6**), native fold_wt_tp63_dbd_tet_mcs97_zn achieved higher DNA-interface confidence (prot_dna_iptm_max = 0.63, prot_dna_pae_min = 4.07 Å, ipTM = 0.85) than palindromic DNA (0.54, 5.92 Å, ipTM = 0.74). EEC mutant C312Y preserved global backbone alignment (pLDDT = 93.25, Cα RMSD = 0.85 Å) yet collapsed tetrahedral Zn^2+^ coordination (max bond 2.57 → 3.97 Å; min angle 99.32° → 15.53°; **Figure 4C**), while R243W reduced prot_dna_iptm_max to 0.58 and widened DNA PAE to 4.93 Å. In phosphorylated IRF6 IAD dimers (ipTM = 0.60, PAE = 2.77 Å), PPS switch mutant S424L weakened subunit docking (ipTM = 0.58, PAE = 3.21 Å) whereas V274I preserved dimerization (ipTM = 0.61, PAE = 2.64 Å); isolated IRF6 DBD dimers showed weak ISRE clamping (ipTM = 0.23, dropping to 0.19 with a 6.20 Å DNA-frame Cα shift in R84C), underscoring trans-stabilization by the phosphorylated IAD dimer. Notably, per-base-pair TP63–DNA contacts in fold_wt_tp63_dbd_tet_mcs97_zn mirrored *MCS9*.*7* loss-of-function profiles in MaveDB (ρ = 0.6048, *p* = 6.52 × 10^−4^) and AlphaGenome observed skin keratinocyte DNASE (ρ = 0.6040, *p* = 6.65 × 10^−4^), peaking at quarter-site bases 344C, 347G, 354C, and 357G (1, 3, 3, and 2 base H-bonds ≤ 3.5 Å; MaveDB −2.07 to −2.40; DNASE −0.85 to

−1.12; **Figure 2D**; **Figure 4A**).

**Figure 2.**
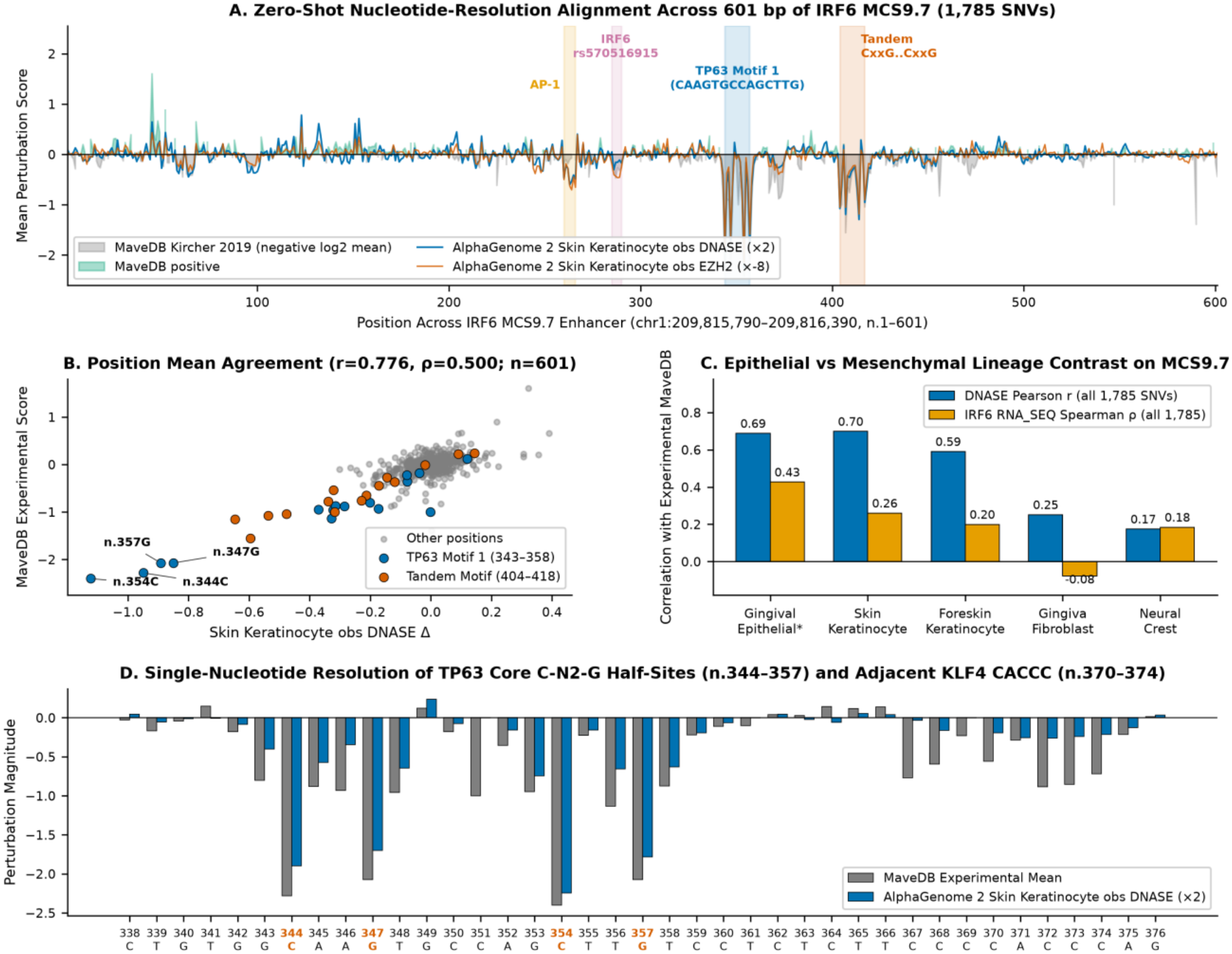
Zero-shot single-nucleotide saturation mutagenesis and lineage specificity across the 601-bp *IRF6 MCS9*.*7* enhancer. (**A**) Nucleotide-resolution alignment across 601 bp of *IRF6 MCS9*.*7* (positions n.1–601; chr1:209,815,790– 209,816,390; *N* = 1,785 SNVs) overlaying experimental MaveDB (Kircher et al., 2019) keratinocyte MPRA scores with AlphaGenome skin keratinocyte observed DNASE (×2) and opposing Polycomb observed EZH2 (×−8) tracks, highlighting AP-1, *IRF6* rs570516915, TP63 Motif 1 (CAAGTGCCAGCTTG), and tandem CxxG..CxxG motifs. (**B**) Position-mean agreement (Pearson *r* = 0.776, Spearman ρ = 0.500; *n* = 601) between MaveDB experimental mean activity and skin keratinocyte observed DNASE Δ, highlighting TP63 Motif 1 (positions 343–358: n.344C, n.347G, n.354C, n.357G) and the Tandem Motif (positions 404–418). (**C**) Epithelial-versus-mesenchymal lineage contrast on *MCS9*.*7*, comparing position-level DNASE Pearson *r* and *IRF6* RNA_SEQ Spearman ρ across Gingival Epithelial *(0*.*69 / 0*.*43;* denotes imputed_exploratory), Skin Keratinocyte (0.70 / 0.26), Foreskin Keratinocyte (0.59 / 0.20), Gingiva Fibroblast (0.25 / −0.08), and Neural Crest (0.17 / 0.18). (**D**) Single-nucleotide resolution across the TP63 core C-N_2_-G half-sites (n.344–357) and adjacent KLF4 CACCC motif (n.370–374) spanning n.338–376, comparing MaveDB position means with AlphaGenome skin keratinocyte observed DNASE (×2) and annotating 344C, 347G, 354C, 357G, and the n.351C *HindIII* cloning outlier.

### AlphaGenome Outperforms Motif Baselines on Clinical Splicing Indels and Prioritizes Cryptic Splice VUS

In Tier NC-1 (*N* = 357 [23 pathogenic, 334 controls] for splicing heads; *N* = 360 for MaxEntScan and RNA_SEQ; **Figure 3A,B**; **Table 3**; **Table S2**), AlphaGenome SPLICE_JUNCTIONS reached AUROC = 0.9876 (AUPRC = 0.9509, Sens@99%Spec = 95.7%), alongside SPLICE_SITES (0.9780) and SPLICE_SITE_USAGE (0.9721), outperforming MaxEntScan (0.9502, Sens@99%Spec = 91.3%), bulk RNA_SEQ (0.9057), and CADD PHRED on its precomputed SNV subset (*N* = 23 [20 pathogenic, 3 controls], AUROC = 0.9083). Donor microdeletion *GRHL3* c.1285+2del scored 0.0 in MaxEntScan and lacked a CADD SNV score yet elicited strong AlphaGenome disruption (SPLICE_SITES = 0.9042, SPLICE_JUNCTIONS = 5.8594), leaving 5′ UTR upstream-AUG allele *IRF6* c.-151G>A (SPLICE_SITES = 0.0078) as the sole missed positive at 99% specificity (**Figure 3B**). Screening 101 ClinVar non-coding VUS (**Table 3**; **Table S1**) prioritized *IRF6* c.174+3A>T (SPLICE_JUNCTIONS = 7.6914), indels c.174+2dup (7.6133) and c.379+4_379+15del (SPLICE_SITES = 0.9577), donor-flanking *TP63* VUS (c.1129+5G>A, c.992+6T>G, c.1746+5G>A), proximal splice-acceptor polypyrimidine-tract VUS *IRF6* c.1061-10C>G (SPLICE_SITES = 0.9667, SPLICE_JUNCTIONS = 5.7656), and *GRHL3* c.1048-1G>T (SPLICE_SITES = 0.9481).

**Figure 3.**
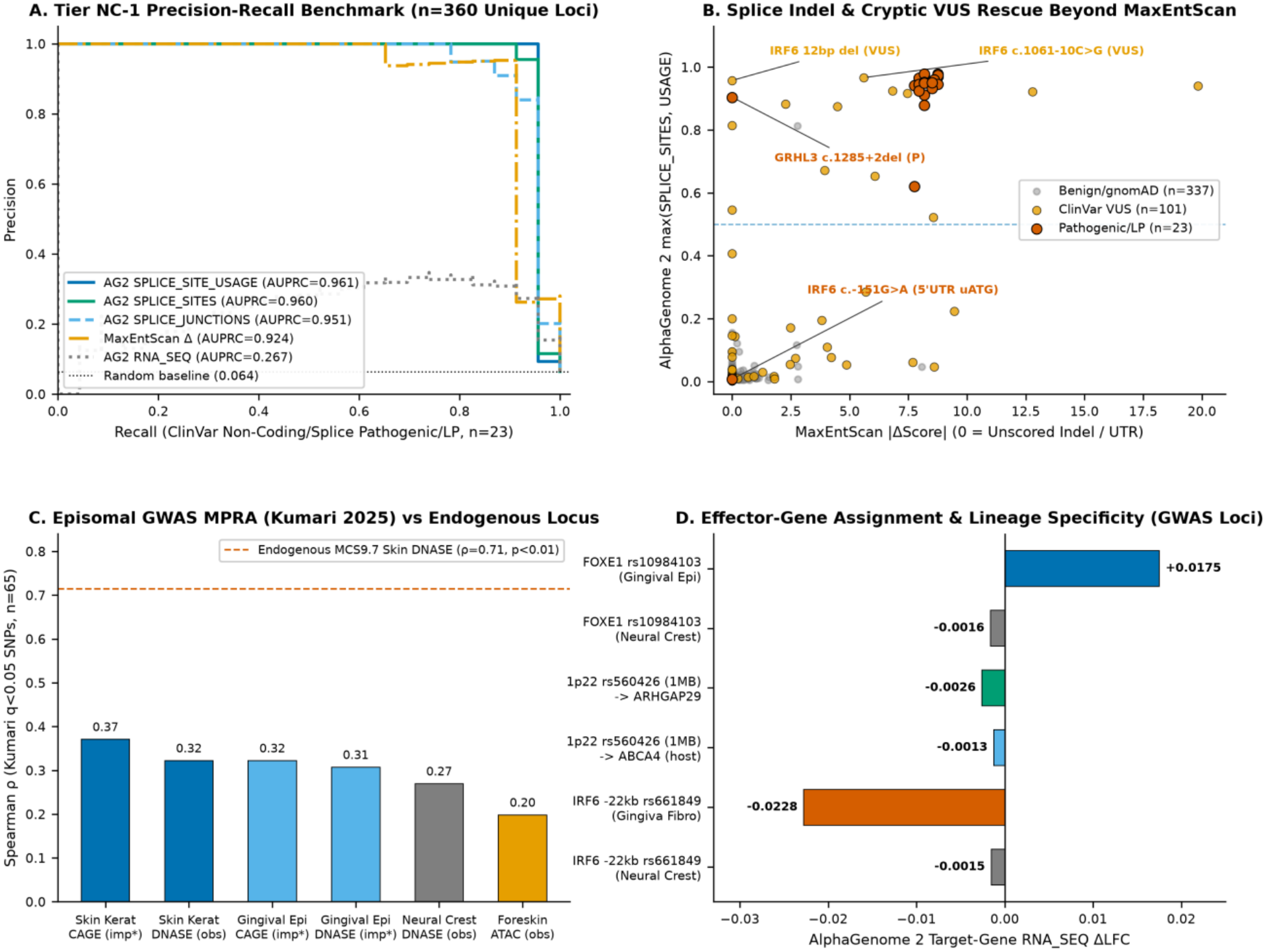
Clinical non-coding splice classification, cryptic splice VUS rescue, and genome-wide OFC MPRA and GWAS effector modeling. (A) Tier NC-1 precision-recall benchmark (*n* = 360 unique loci; *n* = 23 ClinVar pathogenic/likely pathogenic positives) comparing AlphaGenome SPLICE_SITE_USAGE (AUPRC = 0.961), SPLICE_SITES (0.960), SPLICE_JUNCTIONS (0.951), MaxEntScan Δ (0.924), and bulk RNA_SEQ (0.267) against the random baseline (0.064). (B) Splice indel and cryptic VUS rescue beyond MaxEntScan, plotting AlphaGenome max(SPLICE_SITES, USAGE) against MaxEntScan |ΔScore| across pathogenic/likely pathogenic variants (*n* = 23)—highlighting donor deletion *GRHL3* c.1285+2del and 5′ UTR upstream-AUG variant *IRF6* c.-151G>A (uATG)—and ClinVar non-coding VUS (*n* = 101), including *IRF6* 12-bp deletion c.379+4_379+15del and proximal acceptor variant *IRF6* c.1061-10C>G. (**C**) Tier NC-3 zero-shot Spearman rank concordance (ρ) across statistically significant episomal GWAS MPRA variants (Kumari et al., 2025; *q* < 0.05, *n* = 65) for Skin Keratinocyte CAGE (imp; *0*.*37), Skin Keratinocyte ‘DNASE’ (obs; 0*.*32), Gingival Epithelial ‘CAGE’ (imp*; 0.32), Gingival Epithelial DNASE (imp; *0*.*31), Neural Crest ‘DNASE’ (obs; 0*.*27), and Foreskin Keratinocyte ‘ATAC’ (obs; 0*.*20), benchmarked against endogenous* MCS9.7 *Skin ‘DNASE’ (ρ = 0*.*71*, p *< 0*.*01; imp* denotes imputed_exploratory). (**D**) Distal effector-gene assignment and lineage specificity across OFC GWAS loci using AlphaGenome target-gene RNA_SEQ Δlog-fold-change (1-MB window) for *FOXE1* rs10984103 (Gingival Epithelial *+0*.*0175 vs Neural Crest −0*.*0016)*, 1p22 *rs560426 (*ARHGAP29 *−0*.*0026 vs* ABCA4 *−0*.*0013), and* IRF6* −22 kb rs661849 (Gingiva Fibroblast −0.0228 vs Neural Crest −0.0015).

**Figure 4.**
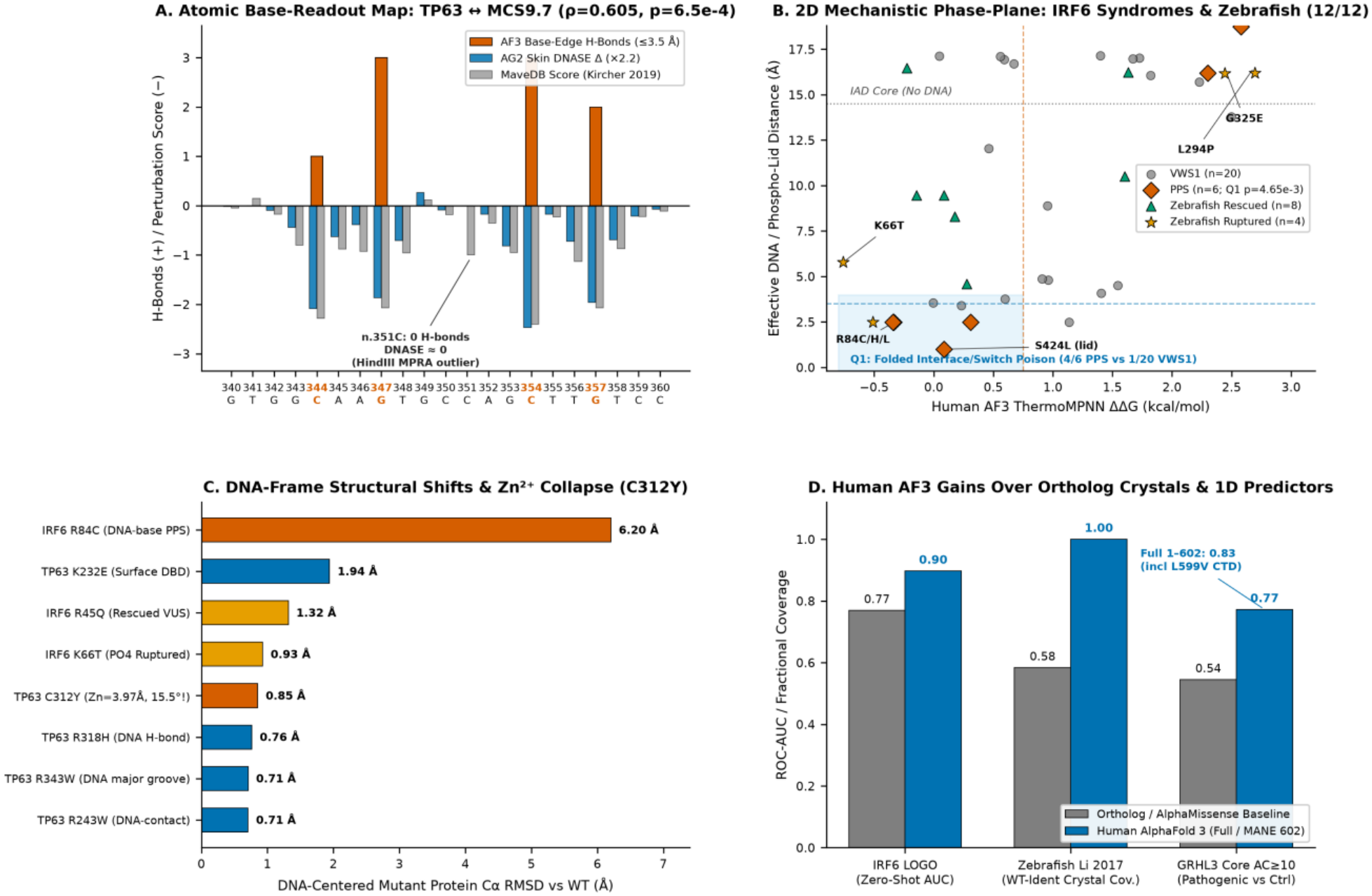
Atomistic protein–DNA base readout, the two-dimensional *IRF6* mechanistic phase plane, and human AlphaFold 3 structural gains. (**A**) Atomic base-readout map of the AF3 TP63 DBD tetramer bound to *MCS9*.*7* DNA (fold_wt_tp63_dbd_tet_mcs97_zn; ρ = 0.605, *p* = 6.5 × 10^−4^), plotting upward base-edge hydrogen bond counts (≤ 3.5 Å: 344C [1], 347G [3], 354C [3], 357G [2]) against downward AlphaGenome Skin DNASE Δ (×2.2) and MaveDB scores across n.340–360, highlighting n.351C (0 H-bonds, DNASE ≈ 0, *HindIII* MPRA cloning outlier). (**B**) Two-dimensional human AF3 mechanistic phase plane for *IRF6* syndromes and zebrafish *in vivo* assays (12/12), plotting ThermoMPNN ΔΔG (kcal/mol) against effective DNA / phospho-lid distance (Å) to isolate Quadrant I folded interface/switch poisons (Q1: 4/6 PPS vs 1/20 VWS1; Fisher’s exact VWS1-versus-PPS OR = 0.0263 [reciprocal PPS enrichment OR = 38.0], *p* = 4.65 × 10^−3^; highlighting R84C/H/L and phospho-lid S424L) alongside lethal-rupture (*n* = 4: K66T, R84C, L294P, G325E) versus mRNA-rescued (*n* = 8) zebrafish alleles. (**C**) DNA-frame structural shifts and Zn^2+^ coordination collapse across mutant AF3 complexes, showing DNA-centered mutant protein Cα RMSD relative to WT (from IRF6 R84C at 6.20 Å to TP63 R243W at 0.71 Å) and highlighting TP63 C312Y (0.85 Å Cα RMSD accompanied by tetrahedral Zn^2+^ collapse to 3.97 Å maximum bond length and 15.5° minimum angle). (**D**) Summary of human AF3 gains over paralog crystal templates and 1D predictors across *IRF6* zero-shot biophysical AUROC (0.77 pairwise [0.78 three-gene LOGO] paralog crystal vs 0.90 human AF3), zebrafish (Li et al., 2017) WT-identical structural coverage (0.58 [7/12] paralog crystal vs 1.00 [12/12] human AF3), and *GRHL3* AC ≥ 10 classification (AlphaMissense 0.54 vs AF3 ordered core 0.77 and full MANE Select 1–602 0.83 including L599V). All structures predicted with AlphaFold Server.

### Epithelial Chromatin Tracks Capture *MCS9*.*7* Enhancer Grammar, Cloning Artifacts, and Long-Range GWAS Effectors

In Tier NC-2 across all *N* = 1,785 SNVs of 601-bp *IRF6 MCS9*.*7* (29), position-mean profiles (*n* = 601) aligned between MaveDB and observed skin keratinocyte DNASE (*r* = 0.776, ρ = 0.500), resolving AP-1, *IRF6* rs570516915, TP63 Motif 1, tandem CxxG..CxxG, and KLF4 CACCC (n.370–374) motifs (**Figure 2A–D**; **Table 3**; **Table S3**). Per-SNV correlations in observed skin (ρ_all_ = 0.4846, ρ_*p*<0.01_ = 0.7141) and foreskin keratinocyte DNASE (0.4811 / 0.6898) far exceeded neural crest (0.2418 / 0.2819) and gingival fibroblast DNASE (0.2892 / 0.3932), rising to 0.5524 / 0.7591 in imputed_exploratory gingival epithelial DNASE (+0.5671 across amplicon interior positions 21–581 vs −0.0051 at ≤ 20-bp plasmid edges in observed skin DNASE). Polycomb observed HIST_H3K27me3 correlated inversely (−0.4459 / −0.6855), and severe LoF classification (*n* = 42) achieved AUROC = 0.9098 in observed skin DNASE (0.9243 for observed EZH2 and imputed JUND). Notably, n.351C>A produced the steepest MaveDB drop (−3.00) while n.351C>G/T were neutral and AlphaGenome predicted ∼0 DNASE change across all three (0 TP63 H-bonds), pinpointing a *de novo* 5′-AAGCTT-3′ *HindIII* site cleaved during MPRA plasmid cloning (29) (**Figure 4A**). Regressing imputed JUND (sk_jund_imp) onto observed skin DNASE (*r* = 0.9433) further isolated accessibility-independent motif loss across the *IRF6* autoregulatory element (n.285–290, residual = −0.0499) harboring clefting allele rs570516915 (16).

In Tier NC-3 across *N* = 65 OFC MPRA hits (*q* < 0.05) (22), zero-shot predictions correlated significantly across observed Skin Keratinocyte DNASE (ρ = +0.3223, *p* = 8.829 × 10^−3^), Neural Crest DNASE (+0.2697, *p* = 2.983 × 10^−2^), and Gingival Epithelial CAGE (+0.2659), reaching +0.3710 (*p* = 2.348 × 10^−3^) in imputed_exploratory Skin Keratinocyte CAGE (**Figure 3C**; **Table 3**; **Table S4**). Across *n* = 13 luciferase SNVs (22), AlphaGenome matched activation at *TP63* rs75436877 and *IRF6* rs661849 (risk-versus-nonrisk MPRA +1.03, luciferase +0.58) while flagging episomal divergence at *FOXE1* rs10984103 (MPRA +1.65 vs luciferase −0.58). Multi-window scans (16 KB–1 MB; **Figure 3D**; **Table S4**) predicted local observed DNase loss at *8q24* rs987525 (−0.4983 at 16 KB, −0.3107 at 1 MB in gingival fibroblasts) while target *MYC* (∼1.20 Mb away) exceeded AlphaGenome’s 1.049-Mb (1,048,576-bp) horizon; concordant DNase (+0.0802) and *FOXE1* RNA_SEQ gain (+0.0189 observed foreskin keratinocyte; +0.0175 imputed_exploratory gingival epithelial vs −0.0016 neural crest) at rs10984103; epithelial suppression at *IRF6* −22 kb rs661849 (observed CAGE −0.1531; fibroblast RNA_SEQ −0.0228); and distal *ARHGAP29* capture at *1p22* rs560426 upon expanding from 100 KB (intronic *ABCA4* only) to 500 KB–1 MB (observed foreskin keratinocyte RNA_SEQ −0.0027; imputed_exploratory gingival epithelial *ARHGAP29* −0.0026 vs *ABCA4* −0.0013).

## Discussion

Uniting atomistic biomolecular complex prediction with sub-megabase sequence-to-function modeling across the *TP63*–*IRF6*–*GRHL3* axis demonstrates that resolving *how* variants perturb macromolecular assemblies or lineage-specific chromatin is essential for craniofacial genetics. Whereas scalar evolutionary predictors such as AlphaMissense excel at binary sorting in canonical domains (23), mechanistic stratification is required to distinguish dominant-negative poisons from haploinsufficient alleles, evaluate full-length MANE Select isoforms, and interpret *in vivo* rescue models.

Three coding principles emerge. First, oligomeric and full-length human AF3 models overcome structural template bottlenecks: tetrameric context unmasks TP63 OD interface VUS (Y402H, E422K), while human AF3 assemblies lift *IRF6* zero-shot biophysical AUROC from 0.7844 to 0.8967 and restore 12/12 zebrafish structural coverage (8, 25). Second, modeling the DNA-bound 602-residue MANE Select GRHL3 homodimer resolves C-terminal L599V omitted from 565-residue AlphaMissense tables and reveals inter-subunit burial at R298H (d_olig_ = 2.65 Å, ΔSASA_int_ = 120.35 Å^2^). Third, a two-dimensional stability-versus-interface phase plane explains the classic clinical split between VWS1 and PPS (2, 9). Because IRF6 operates as a phosphorylation-activated homodimer (9, 34), core-unfolding variants (L294P, G325E, or R84G at the Arg84 hotspot) trigger subunit clearance and haploinsufficient VWS1, whereas stable DNA-contact (R84C/H/L, K66T) or C-terminal phosphoswitch lid (S424L) substitutions poison active dimers in PPS and cause embryonic rupture in zebrafish. Rare variants in *PRKCI* were recently shown to cause Van der Woude syndrome and related peridermopathies (38); although no PRKCI–IRF6 complex was modeled among our 29 AF3 runs, S424L’s selective weakening of phosphorylated IRF6 IAD dimerization (ipTM 0.60 → 0.58; PAE 2.77 → 3.21 Å) without monomer unfolding (+0.0865 kcal/mol) nominates impaired PRKCI-mediated Ser424 phosphorylation as a testable biochemical link across human peridermopathies. Across the coding–regulatory boundary, the AF3 TP63–*MCS9*.*7* tetramer links syndromic DBD mutations (C312Y, R243W) directly to enhancer vulnerability at quarter-site bases 344C, 347G, 354C, and 357G, where MaveDB loss of function coincides with keratinocyte DNase collapse and Polycomb (H3K27me3/EZH2) derepression. Clinically, AlphaGenome splicing heads (AUROC = 0.9876) rescue splice indels missed by MaxEntScan and CADD while prioritizing actionable cryptic splice VUS across *IRF6, TP63*, and *GRHL3*. Concurrently, AlphaGenome serves as an endogenous quality-control filter for episomal MPRAs—discounting 5′-AAGCTT-3′ *HindIII* cloning dropouts at n.351C>A and plasmid-edge noise while linking intronic *1p22* rs560426 to distal *ARHGAP29* (19, 20).

Four limitations merit emphasis. First, AF3 displays static backbone conservatism toward point mutations (26)— retaining high global pLDDT on core unfolders (TP63 I549N, IRF6 L294P) and Zn^2+^-collapsing C312Y—necessitating pairing with ThermoMPNN ΔΔG and local coordination geometry. Second, scarce Mendelian missense positives in *GRHL3* (*P* = 4) yield wide CIs, motivating broader sequencing across underrepresented ancestries (12, 13). Third, because gingival_epithelial DNASE is imputed_exploratory, regulatory inferences should remain anchored to observed keratinocyte and gingival tracks pending primary human palatal periderm atlases. Fourth, AlphaGenome’s 1.049-Mb (1,048,576-bp) context horizon captures *ARHGAP29* at *1p22* yet cannot span ultra-long-range loops beyond ∼0.52 Mb from a centered variant, such as the ∼1.20-Mb *8q24* (rs987525)–*MYC* enhancer axis (21). Together, this multimodal framework provides a quantitative blueprint for mechanistic variant interpretation across craniofacial disorders.

## Declarations

### Ethics approval and consent to participate

Not applicable. This computational study analyzed exclusively publicly available, de-identified human genetic variant repositories (ClinVar, gnomAD, UniProt, MaveDB) and previously published human cohort and embryonic zebrafish summary datasets. No human participants were recruited, no identifiable patient-level records were accessed, and no live vertebrate animal experiments were performed; therefore, institutional review board (IRB) approval, informed consent, and animal care (IACUC) approval were not required.

### Consent for publication

Not applicable.

### Availability of data and materials

All benchmarked ClinVar, gnomAD, MaveDB, and MPRA variant coordinates, AlphaFold 3 structural summary metrics, ThermoMPNN stability calculations, and AlphaGenome multimodal variant effect scores are provided in **Tables 1–3** and **Supplementary Tables S1–S6**. All 29 biomolecular structures were predicted with AlphaFold Server: *“Structure predicted with AlphaFold Server; use is subject to the AlphaFold Server Output Terms of Use”* (non-commercial research use only). AlphaGenome predictions were generated strictly under non-commercial research-only terms. All structural classifications, thermodynamic stability estimates, regulatory predictions, and variant prioritizations reported here are strictly for non-commercial research and hypothesis generation and must not be used for clinical variant interpretation, genetic counseling, or patient management without independent experimental and clinical validation.

### Authors’ contributions

F.M.V. conceived and designed the study, executed the AlphaFold 3 structural modeling, ThermoMPNN biophysical calculations, and AlphaGenome non-coding analyses, and wrote the initial manuscript draft. V.G.S. curated clinical craniofacial genotype–phenotype cohorts, verified syndrome and VUS classifications, and contributed to biological interpretation and manuscript revision. A.N.d.T. supervised statistical benchmarking, audited cross-validation and non-coding track provenance, and critically revised the manuscript. All authors read and approved the final manuscript.

### Supplementary Information

- Table S1: Complete list and multimodal AlphaGenome, MaxEntScan, and CADD scores of prioritized ClinVar cryptic splice variants of uncertain significance across *IRF6, TP63*, and *GRHL3*.
- Table S2: Tier NC-1 clinical non-coding benchmark dataset (*N* = 360) comprising ClinVar pathogenic/likely pathogenic variants, boundary cases, and gnomAD population controls with per-modality predictions.
- Table S3: Tier NC-2 single-base *in silico* saturation mutagenesis matrix (*N* = 1,785 SNVs) across the 601-bp *IRF6 MCS9*.*7* enhancer paired with MaveDB MPRA measurements, accessibility-residualized imputed JUND scores, and n.351 5′-AAGCTT-3′ *HindIII* artifact diagnostics.
- Table S4: Tier NC-3 genome-wide OFC MPRA benchmark (*N* = 65, *q* < 0.05), dual-reporter luciferase validation panel (*n* = 13), and multi-window (16 KB–1 MB) distal GWAS locus profiles stratified by track imputation provenance.
- Table S5: Coding missense variant catalog, ThermoMPNN ΔΔG values, 3D interface distances, AlphaMissense scores, Domainome 1.0 and oligomeric saturation benchmarks, *IRF6* PPS-versus-VWS1 phase-plane classifications, and zebrafish *in vivo* validation results across crystal/paralog PDB and human AlphaFold 3 structures.
- Table S6: Structural confidence, protein–protein and protein–DNA ipTM/PAE metrics, DNA-frame Cα RMSD, and Zn^2+^ coordination geometry across all 29 completed AlphaFold 3 runs (10 WT + 19 Mutants), alongside 28-bp TP63–*MCS9*.*7* atomic base-readout contacts.

